# Influence of switching rule on motor learning

**DOI:** 10.1101/243386

**Authors:** Ken Takiyama, Koutaro Ishii, Takuji Hayshi

## Abstract

Humans and animals can flexibly switch rules to generate appropriate motor commands; for example, actions can be flexibly produced toward a sensory stimulus (e.g., pro-saccade or pro-reaching) or away from a sensory stimulus (e.g., anti-saccade or anti-reaching). Distinct neural activities are related to pro- and anti-movement actions; however, the effects of switching rules on motor learning are unclear. Here, we study the effect of switching rules on motor learning using pro- and anti-arm-reaching movements and a visuomotor rotation task. Although previous results support the perfect availability of learning effects under the same required movements, we show that the learning effects trained in pro-reaching movements are partially rather than perfectly available in anti-reaching movements even under the same required movement direction between those two conditions. The partial transfer is independent of the difference in the visual cue, the cognitive demand, and the actual movement direction between the pro- and anti-reaching movements. We further demonstrate that the availability of learning effects trained with pro-reaching movements is partial not only in anti-reaching movements but in reaching movements with other rules and the availability of learning effects trained with anti-reaching movements is also partial in pro-reaching movements. We thus conclude that the switching rule causes the availability of learning effects to be partial rather than perfect even under same planned movements.

**New & Noteworthy**

Most motor learning experiments supported the involvement of planned movement directions in motor learning; the learning effects trained in a movement direction can be available at movement directions close to the trained one. Here, we show that the availability of motor learning effects is partial rather than perfect even under the same planned movements when rule is switched, which indicates that sports training and rehabilitation should include various situations under the same required motions.

## 1 INTRODUCTION

While kicking a soccer ball toward a friend, recreational soccer players frequently fail to achieve the planned movement. After the ball deviates from the intended movement direction, the players modify their motor commands when performing their next action. Humans and animals have the ability of motor learning to decrease prediction error between planned and actual movements (Shadmehr & Mussa-Ivaldi, 1994; Smith et al., 2006; Takiyama et al., 2015; Takiyama & Sakai, 2016; Hayashi et al., 2016).

The features of motor learning have been clarified based on visually guided and goal-directed arm-reaching movements. After learning reaching movements toward a movement direction, the motor learning effects are partially available for reaching movements toward other movement directions (Thoroughman & Shadmehr, 2000; Donchin et al., 2003). The motor learning effects can differ when the executed actions are the same but the planned movement directions are different (Hirashima & Nozaki, 2012; Sheahan et al., 2016). Based on the transfer of learning effects (a possible measure of the relation between two movement patterns), the relation among movement patterns depends on the differences in the planned movement directions.

Previous studies have relied on the following fixed rule: the same instructed target requires the same planned movements. However, humans and animals can act flexibly toward the same sensory stimulus, e.g., we kick a soccer ball toward a friend who belongs to our team, but we kick the ball 33 away from a friend who is traded to an opposing team. This relation between the sensory stimulus and the response is referred to as the stimulus-response map (S-R map) (Munoz & Everling, 2004). Humans and animals can flexibly switch between congruent (movement direction is toward a sensory stimulus, e.g., pro-saccade and pro-reaching movement) and incongruent (movement direction is away from a sensory stimulus, e.g., anti-saccade and anti-reaching movement) S-R maps. Thus, we can plan the same movements for different targets using various rules, e.g., kicking toward a rightward direction when a friend in our (opposing) team is in the rightward (leftward) direction. Because conventional motor learning experiments have relied on one rule (i.e., the congruent S-R map) and suggested that the planned movement direction determines the motor learning effects, it is unclear whether the same motor learning effect applies when the planned movement is the same but the rule is switched.

Here, we study the effects of switching rules on motor learning using pro- and anti-arm-reaching movements and a visuomotor rotation task (Krakauer et al., 2000). Based on previous behavioral and neurophysiological evidence, the following two possibilities are considered. First, the transfer of learning effects is 100% in the same planned movement direction when a rule is switched. This possibility is based on the following behavioral evidence: motor learning experiments that relied on the congruent S-R map (Hirashima & Nozaki, 2012; Sheahan et al., 2016) and a previous study that investigated the transfer from pro- to anti-saccade in saccadic gain adaptation (Collins et al., 2008; Cotti et al., 2009). Second, the transfer is not 100% when the rule is switched even when the planned movement direction is the same. This possibility is based on the following neural evidence: different neural activities have been observed between pro- and anti-reaching movements (Riehle et al., 1994; Klaes et al., 2011) and between pro- and anti-saccade (Munoz & Everling, 2004) in several brain regions. Certain conventional motor learning theories (e.g., the framework of motor primitive) are inspired by neural activities (Thoroughman & Shadmehr, 2000), and the difference in neural activities may be responsible for the less than 100% transfer of learning effects between pro- and anti-reaching movements.

The current study supports the second possibility as follows: switching rules cause partial availability of motor learning effects even in the same planned movement direction. We show that 1) the transfer of learning effects is partial rather than perfect from pro- to anti-reaching movements, 2) the partial transfer is independent of both visual stimulus and cognitive demand, 3) the transfer of learning effects is also partial from anti- to pro-reaching movements, and 4) the transfer of learning effects is also partial from pro-reaching movements to reaching movements in rules other than anti-reaching movements. We thus suggest that switching rule causes partial availability of learning effects even under the same required movements.

## 2 MATERIALS AND METHODS

### 2.1 Ethics Statement

All participants were informed of the experimental procedures in accordance with the Declaration of Helsinki, and all participants provided written informed consent prior to the initiation of the experiments. All procedures were approved by the ethics committee of the Tokyo University of Agriculture and Technology.

### 2.2 Participants

Forty healthy, right-handed volunteers (aged 18–26 years, nine females) participated in our experiments (10 participants in each of the four experiments).

### 2.3 Experimental Design and Statistical Analysis

The participants were asked to perform 8-cm arm-reaching movements with their right arm while holding the handle of the manipulandum (Geomagic 1.5 HF; Geomagic, Rock Hill, SC, USA). The position of the handle was displayed as a white cursor (6-mm circle) against a black background on a horizontal screen located above their hand. When the visuomotor rotation was introduced, the cursor position deviated from the handle position by multiplying a rotation matrix.

The movement of the handle was constrained to a virtual horizontal plane (10 cm below the screen) that was implemented by a simulated spring (1.0 kN m^-1^) and dumper (0.1 N per (ms^-1^)). A brace was used to reduce unwanted wrist movements. Before each trial, the participants were required to hold the cursor at its center circle (a 10-mm circle). The position and velocity data of the handle were recorded at 500 Hz.

#### Experiment 1

After a 2-s holding time at the center circle (10-mm circle), the color of the center circle changed according to the experimental conditions. Green and red colors indicated pro- and anti-reaching trials, respectively. In the pro-reaching trials, the participants were required to perform arm-reaching movements to move the white cursor toward the visual cue (a 10-mm circle), the color of which was yellow (Figs. 1A, 2A, 4A). In the trials with the anti-reaching movements, the participants were required to perform arm-reaching movements to move the white cursor toward a position opposite to the visual cue (Fig. 1B, 2B, 4B). After an additional 1-s holding time, a yellow visual cue appeared at either the 180° or 0° locations (180° and 0° locations indicated the 9 o’ clock and 3 o’ clock directions, respectively), and a beep sounded to signal the participant to initiate an arm-reaching movement. The participants were required to move the handle at a peak velocity of 470±45 mm s^-1^ (the target velocity was calculated using the minimum-jerk theory with a movement amplitude of 8 cm and a duration of 0.4 s). A warning message appeared on the screen if the movement velocity of the handle rose above (”fast”) or fell below (”slow”) this threshold value. No online correction was allowed. At the end of each trial, the handle was automatically moved back to the starting position by the manipulandum. The participants practiced using the manipulandum and became accustomed to the experimental settings during 98 trials (70 trials for pro-reaching movements and 28 trials for anti-reaching movements).

**Figure 1.**
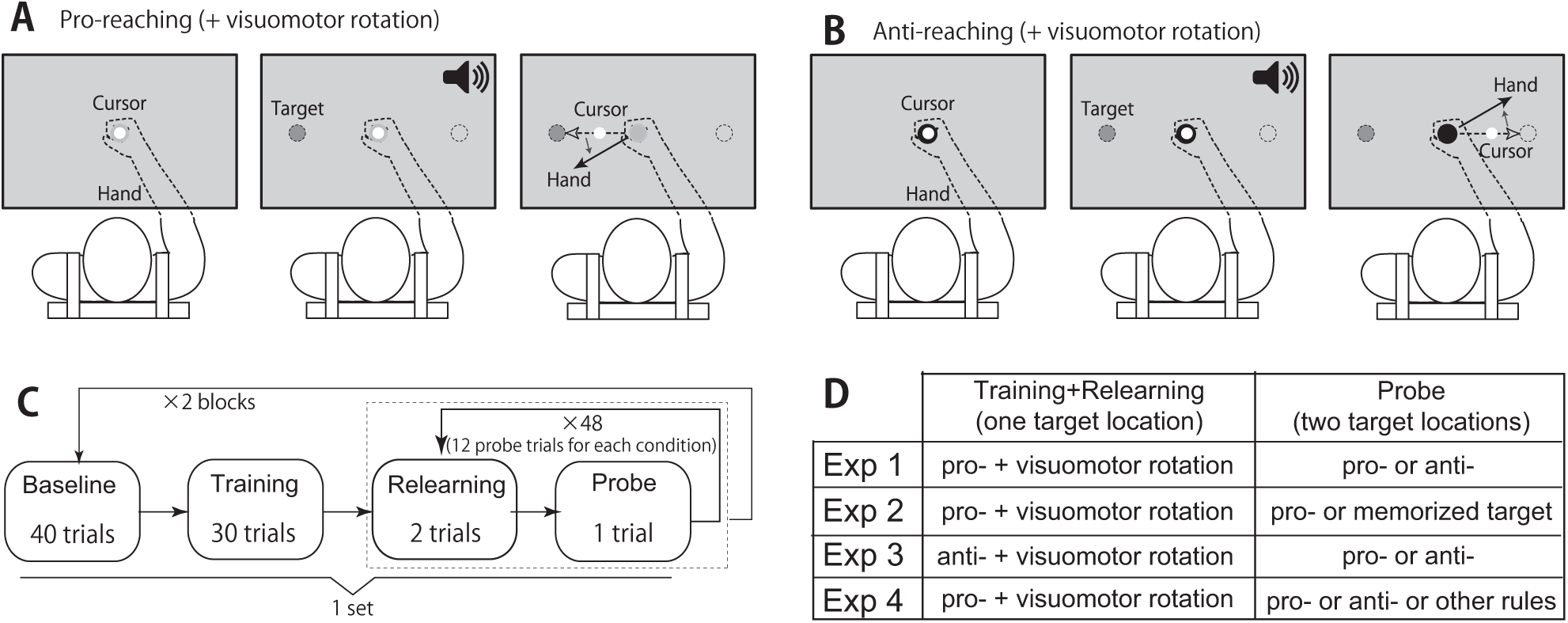
Schematic of the pro- and anti-reaching movements. **A**, Participants performed arm-reaching movements to move the white cursor from the center circle to a target. When the color of the center circle was green (light gray in figure), the participants were required to move the white cursor toward the yellow visual cue (pro-reaching, yellow target was drawn as a dark gray circle). During the training and relearning trials, a visuomotor rotation was imposed. The visuomotor rotation caused the invisible hand movement direction (solid black line) to be deviated from the visible cursor movement direction (dotted black line). **B**, When the color of the center circle was red (black in figure), the participants were required to perform anti-reaching movements to move the white cursor toward the opposite location of the yellow visual cue (dark gray circle in figure). **C**, Experimental protocol for experiments 1, 2, and 4. In experiment 1, participants performed two sets of trials, each of which consisted of 40 baseline trials (20 trials with an invisible cursor trajectory), 30 training trials with a visuomotor rotation and pro-reaching movements, and 48 probe trials with an invisible cursor trajectory. Trials with an invisible cursor trajectory prevented online correction. Before each probe trial, two relearning trials with a visible cursor trajectory, visuomotor rotation, and pro-reaching movements were performed in experiment 1. In experiment 4, pro-reaching movements in training and relearning trials were substituted for anti-reaching movements. In experiment 2, anti-reaching movements in baseline and probe trials were substituted for reaching movements for to-be-memorized targets. **D**, Summary of each experiment.

**Figure 2.**
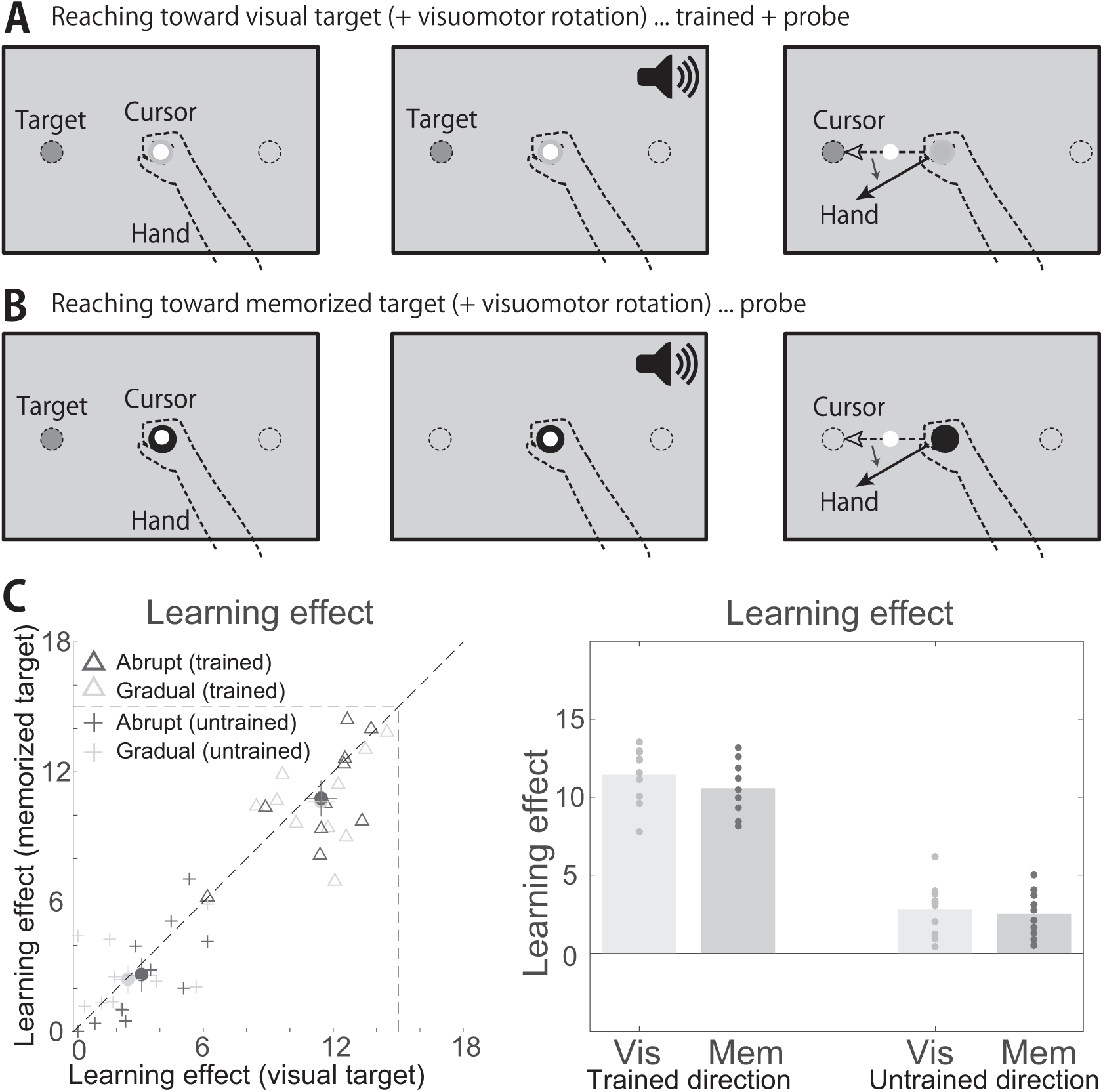
Transfer of learning effects from pro- to anti-reaching movements in experiment 1. **A,B**, Pro- and anti-reaching movements. In experiment 1, visuomotor rotation was introduced in pro-reaching toward single target location and learning effect was probed in both pro- and anti-reaching movements with two target locations. **C**, Learning curves during the training (dotted blue and cyan lines in the former 30 trials), relearning (dotted blue and cyan lines in the latter 96 trials), and probe trials (solid green and red lines for pro- and anti-reaching movements in the probe trials, respectively). Horizontal and vertical axes denote the trial number and learning effect, respectively (the definition of learning effect is provided in the *Materials and Methods* section). Blue and cyan lines denote the learning curves for the visuomotor rotation that was abruptly introduced and gradually increased, respectively. Blue and cyan shaded area denote the standard error of the mean (s.e.m.) (N=10). Green and red error bars also denote the s.e.m. (N=10). **D**, Learning effects in each participant. Open triangles and filled circles denote the learning effects in each participant and mean learning effects in reaching for the trained movement direction. Solid lines crossing the filled circles denote the s.e.m. (N=10). Blue and cyan colors indicate learning effects when the visuomotor rotation was introduced abruptly and gradually increased, respectively. Blue and cyan crosses denote learning effects in reaching for the non-trained movement direction. Open circles and solid lines crossing the circles denote the mean and s.e.m. of the learning effects. There was no significant difference in these learning effects (paired t-test, N=10, p=0.7915 for learning effects toward trained movement direction [open triangles] and p=0.0804 for learning effects toward untrained movement direction [crosses]). **E**, Trajectories of reaching movements and learning effects in pro-(green) and anti-reaching (red) movements during the probe trials. All trajectories were arranged to be directed toward an up-rightward direction. (Left): Trajectories toward the trained movement direction. Thin lines indicate the mean trajectory of each subject during the probe trials in each set. The thick solid lines indicate the mean trajectories across all participants and all sets. Thick dotted lines indicate the mean trajectories in all training trials. (Center left): Averaged learning effects in the trained movement direction across all sets. Each dot denotes the learning effects in each block. Asterisk indicates a significant difference (paired t-test, N=10, p=1.09×10^−5^). (Center right): Trajectories toward the untrained movement direction. (Right): Learning effects in the untrained movement direction. **F**, Relation between the reaction time and learning effect. Reaction time in each block was normalized to satisfy the condition that the mean is zero, and the standard deviation is one. There was no significant difference in the averaged learning effects across the probe trials with the faster and slower reaction times (paired t-test, N=10, p=0.795 for pro- and p=0.977 for anti-reaching movements). **G**, Number of incorrect trials of each participant during the baseline and probe trials when the white cursor was invisible. There was no significant difference in the number of incorrect trials between the pro- and anti-reaching movements (N=20, p=0.900, paired t-test, the combined data from experiments 1 and 3). **H**, Relation between the averaged movement angle in the pro- and anti-reaching movements across all training trials in each set. There was no significant difference in the movement directions between the pro- and anti-reaching movements during the baseline trials (N=80, p=0.967, paired t-test, the combined data from experiments 1 and 3). **I**, Relation between the standard deviation of the movement angle in the pro- and anti-reaching movements during the training trials in each set. There was no significant difference in the standard deviations (N=80, p=0.850, paired t-test).

To eliminate the effect of online correction, the current study probed the learning effects by hiding the white cursor. The baseline movement directions were also calculated by hiding the white cursor before introducing the visuomotor rotation. The learning effect was calculated as 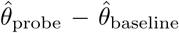, where 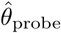 and 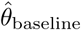 indicated the movement direction after and before adapting to the visuomotor rotation, respectively. 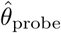 and 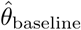 were calculated as 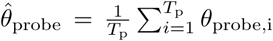 and 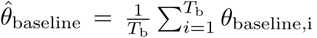, where *T_p_* was the number of successful trials during the probe phase, *θ*_probe,i_ was the movement direction at the *i*th successful trial during the probe phase, *T_b_* was the number of successful trials during the baseline phase, and *θ*_baseline,i_ was the movement direction at the *i*th successful trial during the baseline phase. Because the participants occasionally performed pro-reaching movements during the anti-reaching trials and anti-reaching movements during the pro-reaching movements, *θ*_probe,i_ and *θ*_baseline,i_ were calculated only for the successful trials. We determined the success of each trial according to the movement angle as follows: when the movement angle at all movement times was ±90 within the required movement direction, the trial was considered a success. The movement directions *θ*_probe,i_ and *θ*_baseline,i_ were calculated when the movement velocity reached its peak value.

A visuomotor rotation was applied in the pro-reaching movements, and the transfer of learning effects from the pro- to the anti-reaching movements was investigated in experiment 1. All participants experienced two sets, each of which consisted of 40 baseline trials (target location [either 180° or 0°] and either pro- or anti-reaching movements were pseudorandomly determined [in total, four types of trials]), 30 learning trials (target location was fixed at either 180° or 0° and pro-reaching movements), 96 relearning trials (the fixed target location was the same as that in the learning trials and pro-reaching movements), and 48 probe trials (target location and either pro- or anti-reaching movements were pseudorandomly determined) (Fig. 1C). In 20 of the 40 baseline trials, a white cursor was hidden to calculate the baseline movement direction *θ*_baseline,i_ (i.e., 5 baseline trials for each type of trial). In the subsequent 30 learning trials, the visuomotor rotation was imposed. In the remaining trials, a block of two relearning trials with the white cursor displayed and one probe trial to calculate *θ*_probe,i_, with the white cursor hidden, were repeated 48 times. Because the probe trials cause participants to forget the learning effects, two relearning trials were included to prevent the effects of forgetting. A short break was provided after 20 repetitions (Fig. 1C). Because the short break caused the participants to forget the learning effects, ten learning trials with a 15-degree visuomotor rotation were added after the short break. After one set, 36 washout trials were introduced. The participants experienced a different type of visuomotor rotation in each set, as follows: a 15-degree visuomotor rotation that was abruptly introduced during the learning phase and a visuomotor rotation that increased by 0.5 degrees in each trial, gradually increasing to a maximal value of 15. The order of the two types of visuomotor rotations (abruptly applied or gradually increasing), the direction of the visuomotor rotation (clockwise or counter-clockwise), and the target direction in the learning and relearning trials were counterbalanced across all sets.

#### Experiment 2

In our experimental settings, the anti-reaching movements required arm-reaching movements toward a location without a visual cue (i.e., invisible target). To investigate the effect of reaching toward the invisible target, we conducted experiment 2. In experiment 2, a visuomotor rotation was applied to reaching movements toward a visual cue, and the transfer of the learning effects from those reaching movements to reaching movements toward an invisible and a to-be-memorized target location was investigated (Figs. 3A and B). After a 2-second holding time at the starting position, the color of the center circle changed according to the experimental condition, and a yellow visual cue (a 10-mm circle) appeared at either the 0° or 180° location. When the center circle was red, the target disappeared after 400 ms. Green and red colors indicated reaching movements toward the visual cue and to-be-memorized target, respectively. After an additional 1-s holding time, a beep sounded, signaling the participants to initiate an arm-reaching movement to move the white cursor toward the target. The participants practiced using the manipulandum and became accustomed to the experimental settings over 98 trials (70 trials for reaching movements toward visible targets and 28 trials for reaching movements toward invisible and to-be-memorized targets).

**Figure 3.**
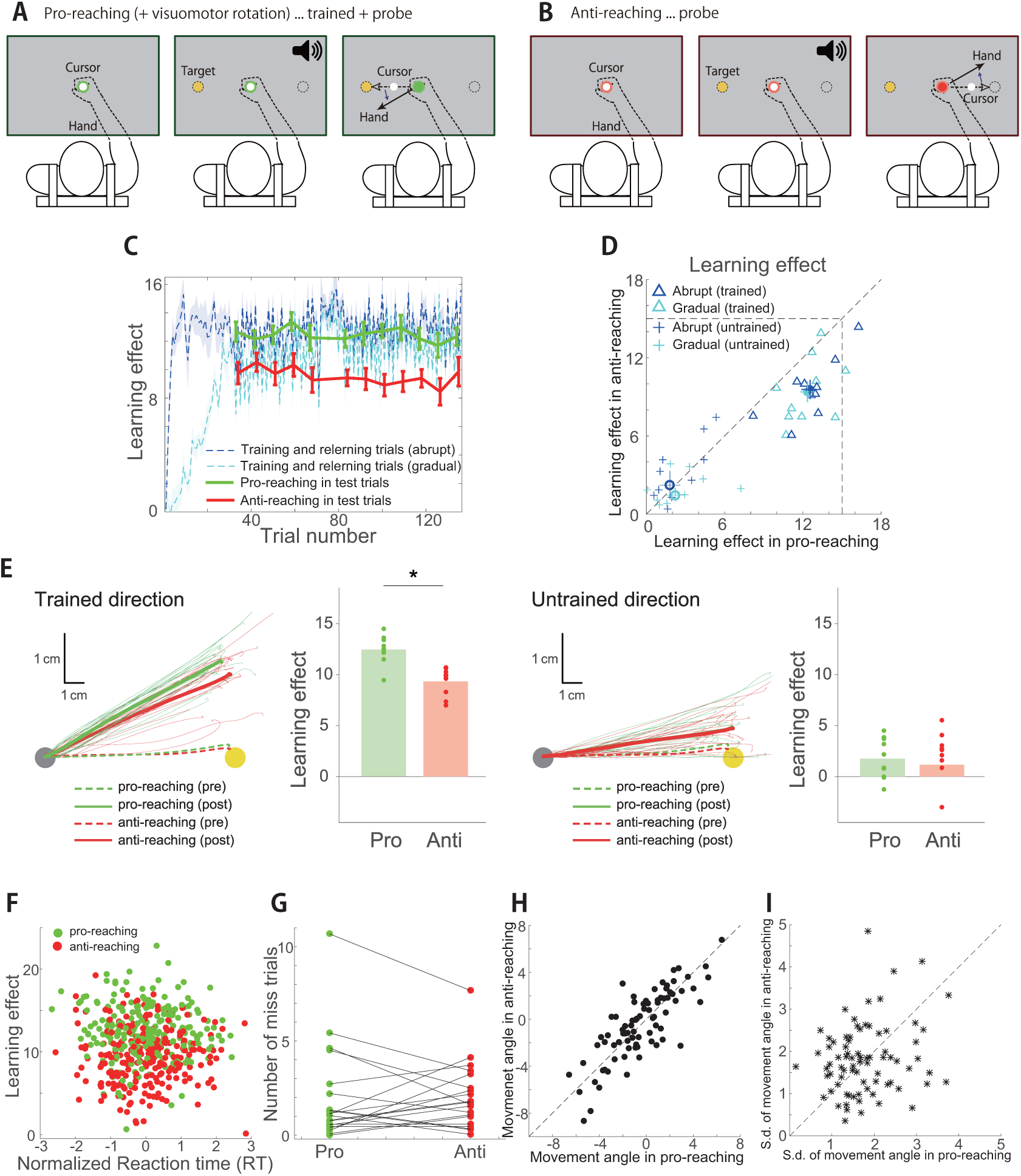
Transfer of learning effects from the reaching movements toward the visible target to those toward the invisible and to-be-memorized targets in experiment 2. **A**, When the color of the center circle was green (light gray in figure), the participants were required to perform arm-reaching movements to move the white cursor toward the yellow visual cue (dark gray in figure). **B**, When the color of the center circle was red (black in figure), the yellow visual cue (dark gray in figure) was visible for 400 ms and subsequently disappeared. The participants were required to memorize the location of the cue and perform arm-reaching movements to move the white cursor toward the memorized cue location after a beep sounded. The beep sounded 600 ms after the disappearance of the cue. **C**, Learning effects in reaching toward visual and to-be-memorized target cues. There was no significant difference between those learning effects in trained movement directions (paired t-test, N=10, p=0.84) and untrained movement directions (paired t-test, N=10, p=0.44).

During the baseline, learning, relearning, and probe trials, the target location and whether the target was to be visible or to-be-memorized were determined following a process similar to that performed in experiment 1, except for that the anti-reaching movements in experiment 1 were substituted for reaching movement toward invisible and to-be-memorized targets in experiment 2.

#### Experiment 3

We conducted experiment 3 to investigate the transfer of the learning effects from anti- to pro-reaching movements. In experiment 3, a visuomotor rotation was applied to the anti-reaching movements, and the transfer of the learning effects from the anti- to the pro-reaching movements was investigated. The experimental settings and protocols were similar to those in experiment 1; however, the learning and relearning trials were performed using anti-reaching movements. One second after a beep sounded, a gray circle appeared at the location where the participants were required to aim (e.g., a gray circle appeared at the leftward location when the visual cue was rightward in the anti-reaching movements). The participants practiced using the manipulandum and became accustomed to the experimental settings over 98 trials (70 trials for the pro-reaching movements and 28 trials for the anti-reaching movements).

During the baseline, learning, relearning, and probe trials, the target location and the pro- or anti-reaching movements were determined using a similar process to that used in experiment 1, except for that the pro-reaching movements in the learning and relearning trials in experiment 1 were substituted for anti-reaching movements in experiment 4.

#### Experiment 4

We conducted experiment 4 to investigate the transfer of the learning effects from pro- to not only anti-reaching movements but also reaching movements with other rules. In experiment 4, a visuomotor rotation was applied to pro-reaching movements, and the transfer of the learning effects from the pro-reaching movements to other reaching movements using the switched rule was investigated. After a 2-second holding time at the center circle (a 10-mm circle), a green or red visual cue appeared at either the 0°, 90°, 180°, or 270° location, and a beep sounded, signaling the participants to initiate the arm-reaching movement (in total, there were eight types of targets). The participants were required to perform 8-cm arm-reaching movements to move the white cursor toward the 180° location when the target color was green (Fig. 5A) and toward the 0° location when the color was red independently of the cue location (Fig. 5B). One second after the beep sound, a gray circle appeared at the location where the participants were required to aim (i.e., a gray circle appeared in the leftward location when the target color was green). The participants practiced using the manipulandum and became accustomed to the experimental settings over 118 trials (78 trials with a yellow target in which the participants were required to perform arm-reaching movements toward the yellow cue and 40 trials in which the participants performed reaching movements toward either the 180° or 0° location depending on the color of the visual cue [green for 180° and red for 0°] that appeared at either 0°, 90°, 180°, or 270°).

**Figure 4.**
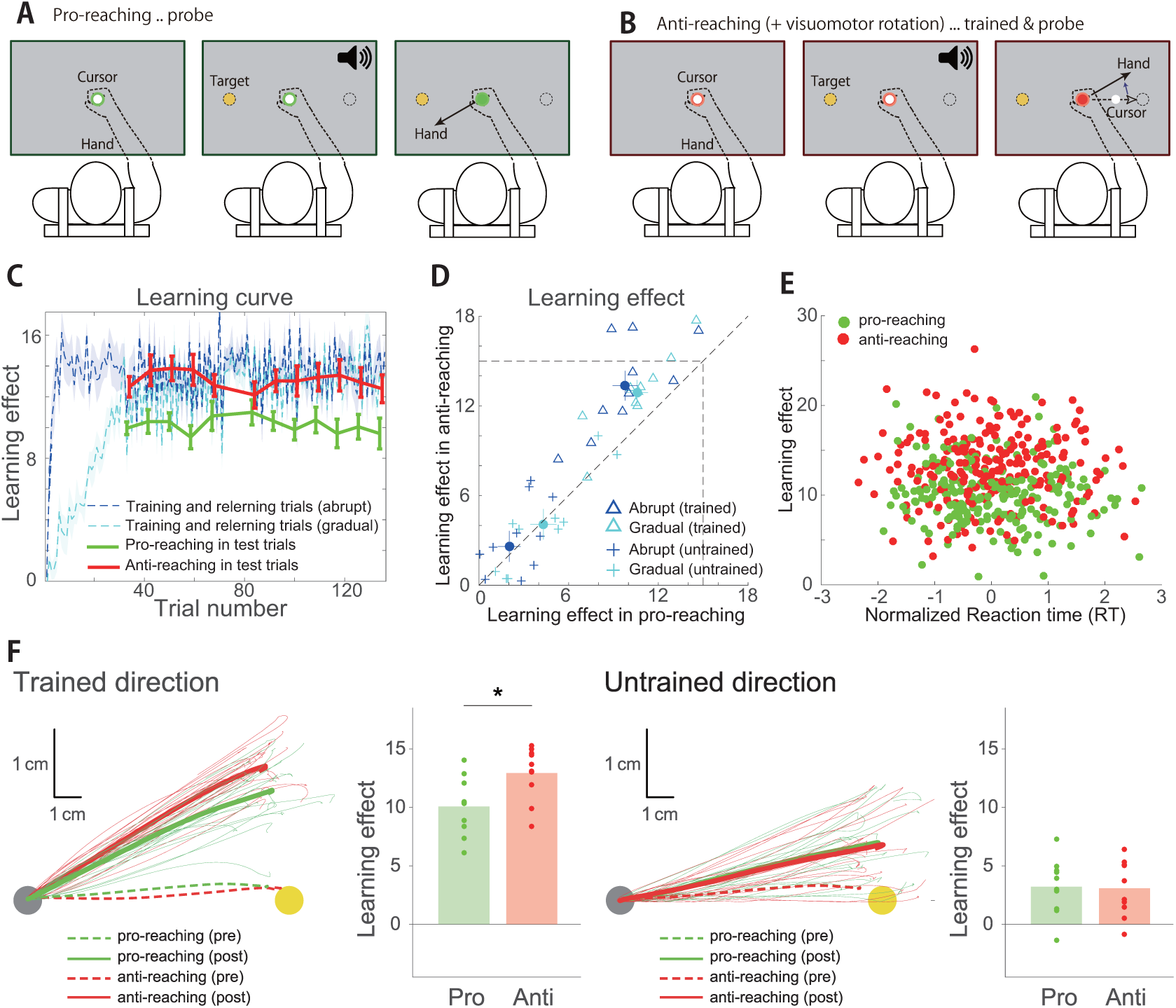
Transfer of learning effects from anti- to pro-reaching movements in experiment 3. **A,B**, Pro- and anti-reaching movements. In experiment 3, visuomotor rotation was introduced in anti-reaching toward single target location and learning effect was probed in both pro- and anti-reaching movements with two target locations. **C**, Learning curves during the training, relearning, and probe trials of the anti-reaching movements. **D**, Learning effects in each subject. **E**, Relation between the reaction time and learning effect. Reaction time in each block was normalized to satisfy the condition that the mean is zero, and the standard deviation is 1. **F**, Trajectories of reaching movements and learning effects. All trajectories were arranged to be directed toward an up-rightward direction. There was significant difference of learning effects in trained movement directions (paired t-test, N=10, p=9.46×10^−4^). In contrast, there was no significant difference of learning effects in untrained movement directions (paired t-test, N=10, p=0.8574).

**Figure 5.**
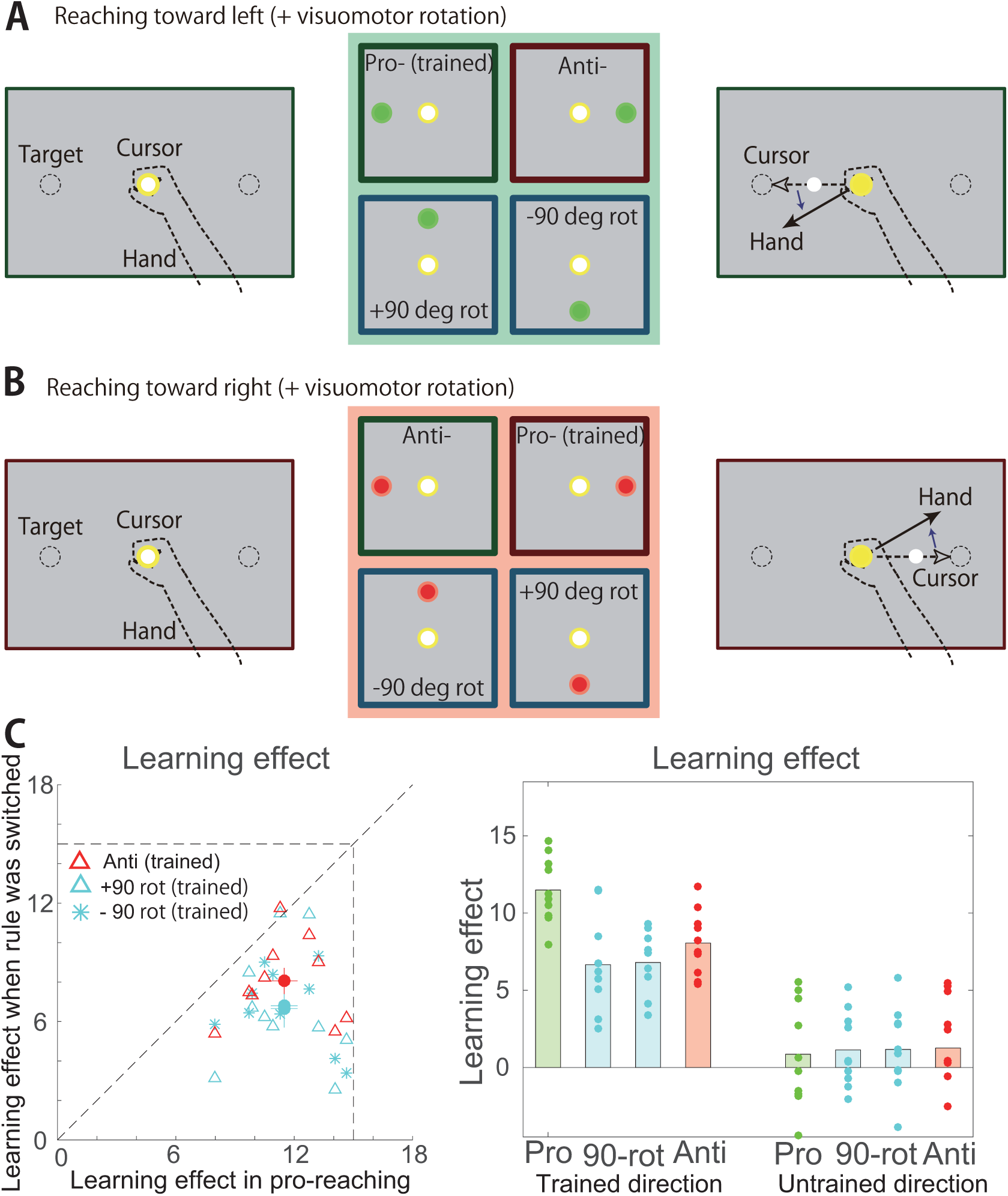
Transfer of learning effects when the rules were switched in experiment 4. **A**, When the color of the visual cue was green, the participants were required to perform reaching movements to move the white cursor toward the 180° direction. **B**, When the color of the visual cue was red, the participants were required to perform reaching movements to move the white cursor toward the 0° direction. **C**, Learning effects in the pro-reaching movements, reaching movements with ±90° rotation, and anti-reaching movements. The learning effects obtained with the pro-reaching movements transferred to other types of reaching movements involving switching rules partially rather than perfectly (paired t-test, N=10, p=0.0029 for reaching movements with a 90 degree rotation, N=10, p=0.0016 for reaching movements with a -90 degree rotation, and N=10, p=0.0047 for anti-reaching movements) in trained movement directions. In untrained directions, there was no significant difference between learning effects in pro-reaching movements and reaching movements with a 90 degree rotation (paired t-test, N=10, p=0.6142), reaching movements with a -90 degree rotation (paired t-test, N=10, p=0.5718), and anti-reaching movements (paired t-test, N=10, p=0.4755).

All participants performed a set of trials that consisted of 80 baseline trials (target color [either green or red] and target location [either 0°, 90°, 180°, or 270°] were pseudorandomly determined), 44 learning trials (target color and location were fixed to either green and 180° or red and 0° for each subject, i.e., pro-reaching movements; Fig. 5A), 192 relearning trials (target color and location were fixed and were the same as those in the learning trials), and 96 probe trials (target color and location were pseudorandomly determined). In 40 of the 80 baseline trials, a white cursor was hidden to calculate the baseline movement direction *θ*_baseline,i_ (i.e., 5 baseline trials for each type of trial). In the following 44 learning trials, a visuomotor rotation was imposed, increasing by 0.5 degrees in each trial to a maximal value of 15 degrees. In the remaining trials, a block of two relearning trials with the white cursor displayed and one probe trial to calculate *θ*_probe,i_ with the white cursor hidden were repeated 96 times.

Two short breaks were provided after 32 blocks. Because the short break caused the participants to forget the learning effects, ten learning trials with a 15-degree visuomotor rotation were added after the short break. In the learning trials, the visuomotor rotation was increased by 0.5 degrees in each trial and reached a maximal value of 15 degrees. The direction of the visuomotor rotation, the target color and the location in the learning and relearning trials were counterbalanced across all participants.

#### Statistical Analysis

We performed paired t-tests.

## 3 RESULTS

### 3.1 Transfer of learning effects from pro- to anti-reaching movements is partial rather than perfect

We evaluated the transfer of the learning effects from the pro- to anti-reaching movements in the same required planned movement direction using a visuomotor rotation of up to 15 degrees. Figure 2C shows the learning curves in the training and relearning trials when either the abruptly applied visuomotor rotation (blue) or gradually increasing visuomotor rotation (cyan) was imposed (used protocols were shown in Fig. 1C). In learning and relearning trials, participants needed to perform pro-reaching movements towards the fixed single target location (either 0° or 180°). Figure 2C also shows the learning effects in the probe trials in the pro- and anti-reaching movements toward a trained movement direction (green and red solid lines indicate the mean±s.e.m., N=10). In probe trials, participants needed to perform either pro- or anti-reaching movements towards either 0° or 180° target location, depending on the color or position of visual cues (in total, there were four types of probe trials; Figs. 2A and 2B). To calculate the learning effects, we removed the incorrect trials (6 out of 120 learning trials were detected to be incorrect).

Figure 2D shows the averaged learning effects of each participant across the successful probe trials when either the abruptly applied visuomotor rotation (blue) or gradually increasing visuomotor rotation (cyan) was imposed in the training and relearning trials. Because there was no significant difference in these learning effects (paired t-test, N=10, p=0.7915 for learning effects toward trained movement direction [open triangles] and p=0.0804 for learning effects toward untrained movement direction [crosses]), we concluded that the type of imposed visuomotor rotation did not affect the transfer of the learning effects from the pro- to the anti-reaching movements, particularly for reaching movements toward a trained movement direction. We thus considered average of those learning effects in each participant for further experimentation.

Figure 2E shows the averaged trajectories and learning effects in each participant. There was a significant difference in the learning effects between the pro- and anti-reaching movements toward the trained-movement direction (paired t-test, N=10, p=1.09×10^−5^, Fig. 2E left and center left panels). The learning effects observed in pro-reaching movements were 12.5±0.4 degrees (mean±s.e.m.), and those in anti-reaching movements were 9.3±0.4 degrees (mean±s.e.m.). The learning effects trained with the pro-reaching movements were thus partially transferred to the anti-reaching movements in the same required planned-movement direction. The movement trajectories also supported this partial transfer. This result indicated that rule switching might affect motor learning.

In addition to rule switching, several factors could explain the partial transfer as follows: 1) the difference in the visual cues between the pro- and anti-reaching tasks affected motor learning; 2) the cognitive demand affected motor learning, 3) the fluctuation of the planned movement direction in the anti-reaching movements affected motor learning, and 4) the reaching movements toward the location without any visual cue (i.e., invisible target) in the anti-reaching tasks affected motor learning. We discuss these possibilities below.

To discuss how the difference in the visual cues affects motor learning (e.g., position of the visual cue), we investigated the transfer of the learning effects from the pro-reaching movements toward a trained movement direction to anti-reaching movements toward an untrained movement direction and the transfer from the pro-reaching movements toward a trained movement direction to the pro-reaching movements toward an untrained movement direction. If the visual cue affected motor learning, the former could be expected to be larger than the latter because the visual cue was congruent in the pro-reaching movements toward the trained direction and anti-reaching movements toward the opposite direction. Our results did not support this hypothesis as follows: there was no significant difference between the two types of transfers (paired t-test, N=10, p=0.653, Fig. 2E center right and right panels).

To investigate the difference in the cognitive demand between the pro- and anti-reaching movements, we calculated the reaction time and the number of incorrect trials. The reaction time in the anti-reaching movements was significantly slower than that in the pro-reaching movements during the probe trials (two-sample t-test, N=240 for pro- and N=234 for anti-reaching movements, p=1.1424×10^−7^, means for pro- and anti-reaching movements were 190.0 and 206.0 ms, respectively), which is consistent with previous studies (Everling & Munoz, 2000; Munoz & Everling, 2004). Because there might be a relation between the learning effects and the reaction time, we calculated the learning effects during the fast and slow reaction time trials in the pro- and anti-reaching movements. There was no significant difference in the averaged learning effects across the probe trials with the faster and slower reaction times (paired t-test, N=10, p=0.795 for pro- and p=0.977 for anti-reaching movements). Figure 2F shows the relation between the learning effects and the normalized reaction times whose mean was zero, and standard deviation was one. There was no significant correlation between the reaction time and learning effects in reaching toward the trained movement direction (N=240, r=-0.0160, p=0.8050 for pro-reaching movements, N=234, r=- 0.0118, p=0.858 for anti-reaching movements). Additionally, we investigated the number of incorrect trials in the pro- and anti-reaching movements during the baseline and probe trials when the cursor disappeared because this number might be indicative of certain aspects of cognitive demand. There was no significant difference in the number of incorrect trials between the pro- and anti-reaching movements (Fig. 2G, N=20, p=0.900, paired t-test). In experiment 1, no incorrect trial was detected in the probe trials with pro-reaching movements, indicating that some incorrect trials with pro-reaching movements occurred during the baseline trials. Because there was no significant difference in the number of incorrect trials between experiments 1 (N=10, p=0.209, paired t-test) and 3 (N=10, p=0.297, paired t-test), the combined data from those experiments are shown in Fig. 2G. We conducted experiment 3 to investigate the transfer of learning effects from anti- to pro-reaching movements, using a similar experimental setting as experiment 1 (see below). Although the reaction time might suggest a larger cognitive demand in the anti- than that in the pro-reaching movements, these results indicated that the cognitive demand might not be significantly relevant to the learning effects in our experimental setting.

Figure 2H demonstrates the difference between the target location and the averaged movement direction of each participant across the baseline trials between the pro- and anti-reaching movements. These values indicate the difference between the actual and required movement directions. There was no significant difference in the movement directions between the pro- and anti-reaching movements during the baseline trials (N=80, p=0.967, paired t-test), which indicated that the fluctuation in the planned movement direction in the anti-reaching movements was not significantly different from that in the pro-reaching movements. Because there was no significant difference in the average movement direction between experiments 1 (N=40, p=0.181, paired t-test, pro- or anti-reaching movements towards 0 or 180 degrees in the first or second block for each participant; in total, four types of reaching movements for each participant) and 3 (N=40, p=0.168, paired t-test), the data from those experiments were combined and are shown in Fig. 2H. Figure 2I demonstrates the standard deviation of the movement angle in the pro- and anti-reaching movements. There was no significant difference in the standard deviations (N=80, p=0.850, paired t-test). Because there was no significant difference in the standard deviation between experiments 1 (N=40, p=0.899, paired t-test) and 3 (N=40, p=0.890, paired t-test), the data from those experiments were combined and are shown in Fig. 2I. These results indicated that the fluctuation of planned movement direction was not highly relevant to the partial transfer in the current experiment.

We conducted experiment 2 to investigate how reaching movements toward an invisible target affected the learning effects. In the pro-reaching movements, the participants were required to reach for visible targets. In contrast, in the anti-reaching movements, the participants were required to reach for invisible targets in the current experimental setting. In addition to the switching rule, the differences between the reaching movements toward visible and invisible targets could affect the transfer of the learning effects. In experiment 2, the participants reached toward either a visible target or invisible and to-be-memorized targets (Figs. 3A and 3B). The participants adapted to a visuomotor rotation in the reaching movements toward a visible target, and the transfer of the learning effects from the reaching movement towards the visible target to those toward the invisible and to-be-memorized targets was investigated. In contrast to the results of experiment 1, the transfer was perfect rather than partial (Fig. 3C, paired t-test, N=10, p=0.578 for reaching movements toward trained movement direction with abruptly applied visuomotor rotation, N=10, p=0.451 with gradually increasing visuomotor rotation, N=10, p=0.841 when combined and averaged). We thus concluded that the reaching movements toward an invisible target were not a significant factor in the partial transfer from the pro- to anti-reaching movements.

### 3.2 Transfer of learning effects from anti- to pro-reaching movements is partial rather than perfect

We further considered the effect of the cognitive demand in the experiment 3, which investigated the transfer of learning effects from anti- to pro-reaching movements. In experiment 3, the participants adapted to the visuomotor rotation in anti-reaching movements. We then investigated the transfer from anti- to pro-reaching movements. Our results indicated that the transfer from anti- to pro-reaching movements was partial rather than perfect (Fig. 4, paired t-test, N=10, p=9.46×10^−4^ for reaching movements toward the trained movement direction with abruptly introduced visuomotor rotation, N=10, p=1.68×10^−4^ for reaching movements toward the trained movement direction with gradually increasing visuomotor rotation, and N=10, p=4.21×10^−4^ when combined). Additionally, there was no significant difference between the degree of transfer from the pro- to anti-reaching (0.748±0.026 [mean±s.e.m.]) and from the anti- to pro-reaching movements (0.778±0.039 [mean±s.e.m.], two-sample t-test, N=10, p=0.529).

In experiment 3, the participants experienced more anti-reaching movement trials than pro-reaching movement trials. The reaction time could be expected to be the same or faster in the anti-reaching movements than that in the pro-reaching movements; however, the reaction time was still slower in the anti-reaching movements than that in the pro-reaching movements during the probe trials (two-sample t-test, N=232 and N=238 for pro- and anti-reaching, respectively, p=4.1685×10^−4^, means for pro- and anti-reaching movements were 193.6 ms and 206.8 ms, respectively), indicating that the cognitive demand might be larger in the anti-reaching movements. If cognitive demand affected motor learning, the transfer from anti- to pro-reaching movements should be larger than 100% or close to 100%. Because we could not find this tendency, cognitive demand was not considered a significant factor affecting motor learning in switching rules in the current experimental setting.

### 3.3 Switching rules caused partial availability of motor learning

To investigate whether switching rules affects motor learning further and whether anti-reaching is a specific factor that affects motor learning, we conducted experiment 4 (Fig. 5). In this experiment, the participants were required to reach for the 0° location when the target color was green independently of the location of the visual cue. When the color was red, the participants were required to reach for the 180° location independently of the location of the visual cue. This experimental setting included not only anti-reaching but also other types of rules (i.e., the color of the visual cue was a determinant of the required movement direction) (Figs. 5A and 5B). Subjects adapted to the visuomotor rotation with pro-reaching movements. We then investigated the transfer of the learning effects from the pro-reaching to other types of reaching movements. The learning effects obtained with the pro-reaching movements transferred to other types of reaching movements involving switching rules partially rather than perfectly (Fig. 5C, paired t-test, N=10, p=0.0029 for reaching movements with a 90 degree rotation, N=10, p=0.0016 for reaching movements with a -90 degree rotation, and N=10, p=0.0047 for anti-reaching movements). There was a significant difference between the learning effects in the reaching movements with a 90 degree rotation and those in the anti-reaching movements (paired t-test, N=10, p=0.0278), although there was no significant difference in the learning effects between the reaching movements with a -90 degree rotation and those in the anti-reaching movements (paired t-test, N=10, p=0.0658) and those with a 90 degree rotation and -90 degree rotation (paired t-test, N=10, p=0.8873). The partial transfer from the pro-reaching to other types of reaching indicated that anti-reaching was not a specific factor that induces a partial transfer; therefore, the switching rules affected motor learning and caused partial availability of learning effects.

## 4 DISCUSSION

The current study investigated the influence of switching rules on motor learning in the same required planned movement, and we observed that switching rules decreased the learning effects. A decrease in learning effects was observed when the transfer of learning effects from pro- to anti-reaching movements (Fig. 2), from anti- to pro-reaching movements (Fig. 4), and from pro-reaching to reaching movements under several types of rules was investigated (Fig. 5). The decrease in learning effects might not be attributed to the reaching movements toward the location without any visual cue (Fig. 3), cognitive demand (Figs. 2F, 2G, and 4E), and fluctuation of aiming direction (Figs. 2H and 2I). Our results suggested that switching rule causes partial availability of learning effects even under the same planned movements.

### 4.1 Partial availability of learning effects in the same planned movements with different rules

Previous studies reported that a principal factor in determining the availability of motor learning effects is the planned movement direction (Thoroughman & Shadmehr, 2000; Donchin et al., 2003, Hirashima & Nozaki, 2012; Sheahan et al., 2016). After the training, the learning effects are partially available in the probe trials when the movement directions are different between the trained and probed movement directions (Thoroughman & Shadmehr, 2000; Donchin et al., 2003). Even when the executed movement directions are the same, the participants can learn different motor tasks when the planned movement directions are different (Hirashima & Nozaki, 2012; Sheahan et al., 2016). Based on these results, we expected the same amount of motor learning effects in the pro- and anti-reaching movements when the same movement directions were required; however, the current study revealed that the availability of motor learning effects was partial rather than perfect when the same planned movement directions were needed, but the rule was switched (Figs. 2, 4, and 5). Our results indicate that the partial availability originated from the modulation of neural activities when the same planned movement directions were required, but the rule was switched. The modulation was observed in the pro- and anti-saccade, pro- and anti-reaching, and several brain regions related to saccade or arm-reaching movements (Riehle et al., 1994; Klaes et al., 2011, Munoz & Everling, 2004, Everling & Munoz, 2000, Schlag-Rey et al., 1997).

### 4.2 Influence of cognitive demand on motor learning

Previous studies have reported that divided attention affects motor learning (Taylor & Thoroughman, 2007); more required attention correlates with less motor adaptation. Anti-reaching movements might require higher attention levels than pro-reaching movements because the reaction time was significantly slower in the anti-reaching movements. If attention or cognitive demand affected motor learning in the current situation, the learning effects would be larger in the trials with either faster or slower reaction times, the transfer from the anti- to the pro-reaching movements would be an over-generalization (transfer rate larger than one), or the number of incorrect trials would be larger in the anti-reaching movements than that in the pro-reaching movements. Although there may be a difference in the attention requirement between the pro- and anti-reaching movements, we could not find a significant effect of the difference in the current study (Fig. 2). These results indicate that the influence of the modulation of neural activities on motor learning overrides that of divided attention.

### 4.3 Influence of explicit and implicit learning

Learning effects consist of (at least) two factors; explicit (cognitive) and implicit learning effects (Taylor et al., 2014; Morehead et al., 2015; Butcher et al., 2017). In a visuomotor rotation, participants can compensate for task error between the target position and actual movement by aiming at a different location from the target position. This cognitive factor to change aiming direction is referred to as explicit learning, which is distinct from implicit learning in which movement is changed to minimize prediction error without awareness. In our experiments, there was a possibility that subjects relied on explicit learning in the trained condition while these explicit learning effects were unavailable in the probe trials, which would imply that the availability of learning effects in probe trials was partial rather than perfect. Possible arguments against this possibility are the following; 1) visuomotor rotation of 15 degrees, 2), short preparation time, and 3) unawareness of the perturbation. First, we used the visuomotor rotation of 15 degrees because explicit learning effects for adapting to a visuomotor rotation of 15 degrees were likely less than those for a visuomotor rotation of 30 degrees (Fig. 4 in Morehead et al., 2015, Fig. 3 in Butcher et al., 2017). Second, the movement preparation time was quite short in our experimental setting, as subjects needed to move the cursor towards the target immediately after it appeared, without any delay. Because a short preparation time can suppress explicit learning (Leow et al., 2017), our results mainly reflect implicit learning effects. Third, no participant was aware of the visuomotor rotation even when the visuomotor rotation of 15 degrees was applied abruptly (N = 40). After each experiment, we asked the participants ”Please tell us what you were aware of in this experiment.” No participants reported the existence of visuomotor rotation. Because unaware perturbation cannot be explicitly compensated for, our results mainly reflect implicit learning effects.

### 4.4 Relation between arm-reaching movement and saccade

A previous study investigated the transfer of learning effects from pro- to anti-saccade using saccadic gain adaptation (Collins et al., 2008; Cotti et al., 2009). The difference between saccadic gain in prelearning and that in post-learning was the same as that between pro- and anti-saccade, which suggested the perfect transfer from pro- to anti-saccade. Although this result may be inconsistent with the current results, several factors could potentially explain this difference. First, the difference in the effector for the required movements may be an explanatory factor; the previous study relied on saccade, and the current study relied on arm-reaching movements. Because certain brain regions have different activities for saccade and arm-reaching movements (Snyder vet al., 2000; Cui et al., 2007), the difference in the effector can affect the transfer of the learning effects. Furthermore, the difference in the adaptation task may be an explanatory factor. The previous study relied on gain adaptation or an adaptation of the movement distance. In contrast, the current study focused on adaptation to the visuomotor rotation because previous studies have focused on adaptation to the angular deviation or deviation perpendicular to the movement direction more than movement amplitude (Thoroughman & Shadmehr, 2000; Donchin et al., 2003; Taylor et al., 2014; Takiyama et al., 2015; Hayashi et al., 2016). Several previous studies have reported that the transfer of learning effects from a trained movement to an untrained (probed) movement was different between gain adaptation and the adaptation to the visuomotor rotation (Krakauer et al., 2000), indicating that the neural mechanisms underlying these adaptations are different. The difference in both the effector and adaptation task could contribute to the difference between the results of the previous study (Collins et al., 2008; Cotti et al., 2009) and the current results.

## 5 ACKNOWLEDGMENTS

We thank D. Nozaki and S. Kasuga for their helpful comments.

## Author contributions

K.T. and K.I. designed and performed the experiments. K.T. and T.H. performed the analyses and wrote the manuscript.

